# The Highly Selective 5-HT2B Receptor Antagonist MW073 Mitigates Aggressive Behavior in an Alzheimer’s Disease Mouse Model

**DOI:** 10.1101/2025.09.21.677617

**Authors:** Erica Acquarone, Saktimayee M. Roy, Agnieszka Staniszewski, Martin D. Watterson, Ottavio Arancio

**Affiliations:** Taub Institute for Research on Alzheimer’s Disease and the Aging Brain, 630 West 168th Street, P&S 12-420D, New York, NY, 10032, USA; Department of Pharmacology, Feinberg School of Medicine, Northwestern University, 320 E. Superior St. Chicago, IL, 60611, USA; Department of Pathology and Cell Biology, Columbia University, New York, NY, 10032, USA; Department of Medicine, Columbia University, New York, NY, 10032, USA

## Abstract

Alzheimer’s disease (AD) is a multifactorial neurodegenerative disorder and the leading cause of dementia worldwide. Progressive synaptic and neuronal loss underlies the decline in cognition and daily functioning, often accompanied by behavioral and psychological symptoms. Among these, neuropsychiatric disturbances such as agitation and aggression affect 20–65% of patients and represent a major source of caregiver burden. The serotonin receptor antagonist MW073 has recently emerged as a potential therapeutic candidate, showing efficacy in counteracting Aβ- and tau-induced synaptic and memory deficits in AD mouse models. Here, we investigated whether MW073 also mitigates aggressive behavior in Tg2576 mice, a widely used AD model that also displays heightened aggressiveness. Our findings demonstrate that MW073 significantly reduced aggressive tendencies in Tg2576 mice, suggesting that serotonergic modulation may represent a promising strategy to address both cognitive and neuropsychiatric symptoms of AD.

## 2. INTRODUCTION

Alzheimer’s disease (AD) is a multifactorial neurodegenerative disease and the most common form of dementia. AD is characterized by the abnormal aggregation and deposition of amyloid-β peptides into extracellular plaques and hyperphosphorylated tau protein into intracellular neurofibrillary tangles. (Zhang et al 2024) These two neuropathological hallmarks of the disease are followed by synaptic and neuronal loss resulting in progressive cognitive and functional decline.(Chatzikostopoulos et al 2025) The etiology od AD is complex and the precise mechanisms underlying its onset are not yet completely understood. There is a spectrum of other factors that can contribute to the pathology of AD, such as cholinergic deficit, neuroinflammation, glutamate deficit, oxidative stress, insulin resistance mitochondrial dysfunction and deficit in autophagy. (Beata et al 2023, Stanciu et al 2019, Thakral et al 2023)

With the progression of the disease, there is another aspect that is important and becomes progressively more prevalent: the neuropsychiatric symptoms (NPS) (Lyketsos et al 2011). Aggressivity, apathy, agitation, and psychosis are the most common NPS. (Kales et al 2014, Yu et al 2019). There are not treatments that have been tailored to specifically alleviate AD patient NPS symptoms (Ijaopo 2017). In the early 2000s, in Canada and Australia, an atypical antipsychotic drug named Risperidone was used for the short-term treatment of persistent and severe aggression in AD. It has been widely used off-label to treat behavioral and psychological symptoms of dementia (BPSD), including agitation, aggression, and psychosis. (Yunusa & El Helou 2020) In the past decade, safety concerns emerged regarding risperidone due to increased risk for cerebrovascular adverse events and death in the elderly population. The drug was never approved in the United States. Nowadays, clinical guidelines suggest that pharmacological treatments might be considered when nonpharmacological treatments fail and just in severe form of dementia in which the patient is threatening to harm himself or others. In 2023, in the United States, the Food and Drug Administration (FDA) approved Brexpiprazole, a newer atypical antipsychotic, for the treatment of agitation in AD (https://www.fda.gov/news-events/press-announcements/fda-approves-first-drug-treat-agitation-symptoms-associated-dementia-due-alzheimers-disease).

Serotonin (5-hydroxytryptamine, 5-HT) is a monoamine neurotransmitter derived from the amino acid tryptophan. It plays a pivotal role in regulating a wide array of physiological and behavioral processes, including mood, aggressivity, appetite, sleep, cognition, pain perception, thermoregulation, cardiovascular function, and gastrointestinal motility (Berger et al 2009). MW073 is a highly selective antagonist of the serotonin 5-HT2b receptor (5-HT2bR) (Roy et al 2025). It demonstrates concentration-dependent inhibition of receptor activity with no detectable agonism at any of the 165 GPCRs tested and no significant off-target activity in kinome, transporter, off-target enzyme, or drug target ion channel screens. Precision selectivity positions MW073 as a first in-class, pharmacologically derisked probe for 5-HT2bR inhibition.

MW073 molecular mechanism of action is dose dependent inhibition of serotonin activation of 5-HT2bR and its β-arrestin-1 recruitment in both human and mouse assays. The prevailing view is that the cellular signaling mechanism would be attenuation of the Gq → PLC-β2 → DAG/IP_3_ signaling cascade and downstream calcium signaling, with potential protection from synaptic dysfunction. Inhibition of arresting recruitment could potentially attenuate receptor internalization and associated signaling. A MW073 mechanism of action that integrates competitive antagonism and β-arrestin inhibition via serotonergic signaling offers the potential to preserves synaptic function and restore behavioral outcomes in disease models. Consistent with the hypothesis, treatment of mouse models with MW073 preserved synaptic function in the presence of the AD relevant stressors Aβ and tau oligomers. The selectivity of the efficacy was documented by no effects on locomotion, anxiety, motivation, or sensory thresholds. Taken in their entirety, the data establishes a potential new precision pharmacological pathway for treating AD associated cognitive dysfunction.

MW073 also represent a uniquely potent, selective, and clean reference standard for probing 5-HT2bR pharmacology in widely used clinical drugs. Using MW073 as a reference standard to test approved neurotropic drugs for potential 5-HT2bR inhibition identified risperidone was found to mimic MW073 as a dose dependent inhibitor of 5-HT2bR activity and β-arrestin-1 recruitment in both human and mouse assays. Risperidone is considered a pleiotropic antipsychotic with multiple dopamine and serotonin receptor activities. The outcomes raised the hypothesis that MW073 might provide a precision pharmacological pathway for treating AD associated neuropsychiatric syndromes. To probe test this hypothesis, we evaluated MW073 treatment of aggressivity in an established and widely used AD relevant animal model, the Tg2576 mouse. Our objective was to evaluate whether chronic treatment with the antagonist could reduce aggressive behavior, thereby providing preclinical evidence for its potential therapeutic application in managing behavioral disturbances in AD.

## 3. RESULTS

### 3.1 Tg2576 males are more aggressive than Tg females

Our first objective was to evaluate potential sex differences in aggressive behavior in Tg2576 mice. In the resident–intruder paradigm, no Tg2576 resident females (0/16) initiated attacks against intruder females during the 10-minute testing period. In contrast, 13 out of 16 Tg2576 resident males exhibited overt aggression toward intruder males (Fig. 1A–B). Quantitative analysis revealed that both the frequency of attacks and the cumulative attack duration were significantly higher in Tg2576 males compared to females (p < 0.001; Fig. 1A–B). Given the pronounced male-specific aggressive phenotype, all subsequent pharmacological experiments with MW073 were conducted in Tg2576 male residents.

**Fig. 1.**
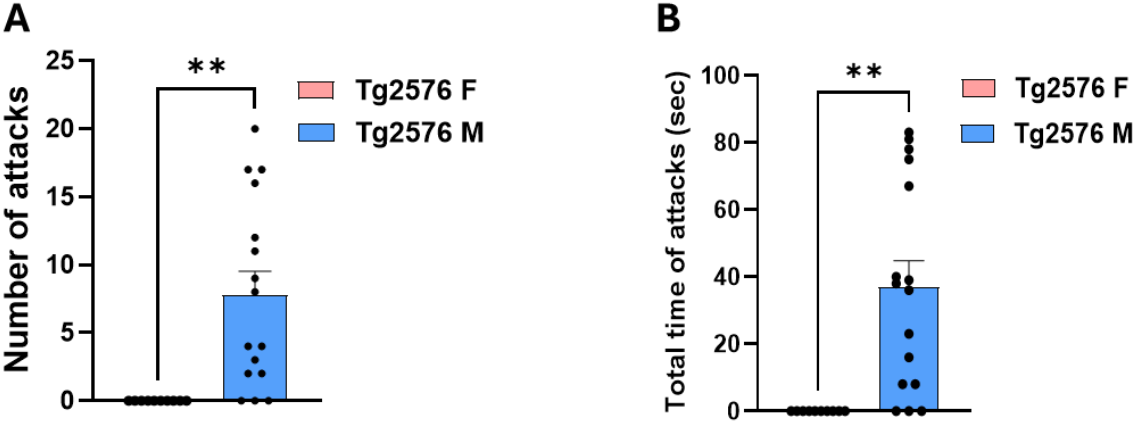
Tg2576 males are more aggressive than Tg2576 females. Bar graphs representing rate scores related to aggressive behavior in Tg2576 resident males and Tg2576 resident females. (**A-B**) The number of attacks, as well as the total attack time, of male Tg2576 animals was dramatically higher than in females Tg2576 mice (number of attacks: Tg2576 females = 0, Tg2576 males = 7.81 ± 1.73, p = 0.0016; total attack time: Tg2576 females = 0 sec, Tg2576 males = 37 ± 7.77 sec, p = 0.0010). These experiments were performed on 10 Tg2576 female mice and 16 Tg2576 male mice. The average age of females was 262.3 ± 29.21, whereas the average age of males was 294.56 ± 34.55 p = 0.3482).

### 3.2 Aggressive behavior test in Tg2576 and non-transgenic (nTg) littermate males

Our next goal was to compare aggressiveness in Tg2576 mice compared to non-transgenic (nTg) males. We found that only 3 of 20 nTg resident males exhibited aggression toward intruder males during the 10-minute resident–intruder test, whereas 11 of 14 Tg2576 residents initiated attacks (Fig. 2A–D). The latency to the first attack was markedly reduced in Tg2576 mice, which began aggression at approximately 2 minutes following intruder introduction, compared to ∼7 minutes in nTg controls (p < 0.01; Fig. 2A). The duration of the initial attack was slightly longer in Tg2576 mice relative to nTg, but this difference did not reach statistical significance (Fig. 2B). By contrast, both the total number of attacks and the cumulative attack duration were significantly elevated in Tg2576 residents compared with nTg littermates (p < 0.001; Fig. 2C–D).

**Fig. 2.**
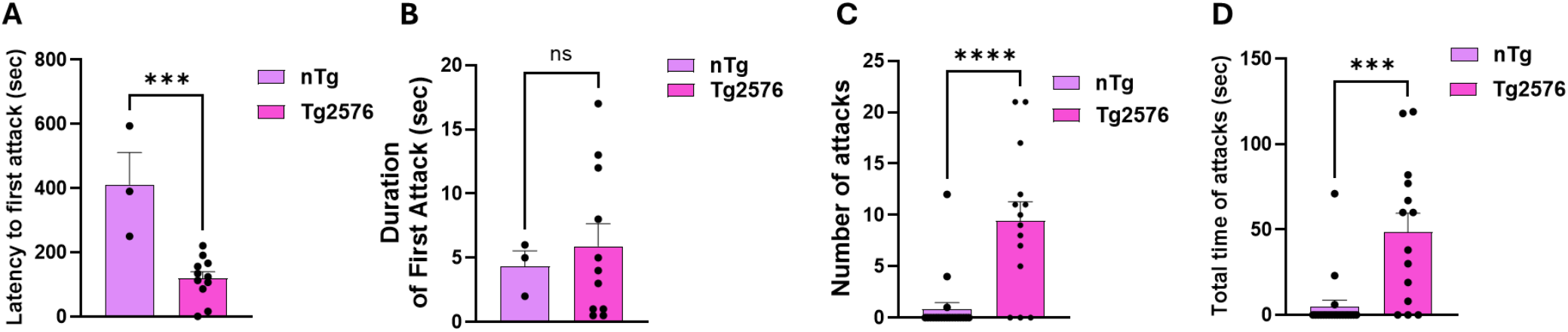
Tg2576 resident males are more aggressive than nTg resident males. Bar graphs representing rate scores related to aggressive behavior in Tg2576 and non-transgenic (nTg) resident males. (**A**) nTg residents displayed longer latency to the first attack compared to Tg2576 mice (nTg = 411.33 ± 99.87 sec, Tg = 119.54 ± 20.27 sec, t-test for this and all the other graphs p = 0.0004) (**B**) Compared to nTg residents, Tg2576 residents showed slightly higher not statistically significant duration of the first attack (nTg = 4.33 ± 1.20 sec, Tg = 5.909 ± 1.74 sec, p = 0.6590). (**C-D**) The number of attacks, as well as the total attack time, of Tg animals was dramatically higher compared to nTg mice (number of attacks: nTg = 0.85 ± 0.62, Tg = 9.429 ± 1.86, p < 0.0001; total attack time: nTg = 5.00 ± 9.46 sec, Tg = 48.43 ± 11.08 sec, p = 0.0002). These experiments were performed on 20 nTg mice and 14 Tg2576 male mice. The average age of nTg mice was 224.2 ± 40.218, whereas the average age of the Tg2576 was 181 ± 13.284 (p = 0.3897). Standard errors and results from individual experiments are shown within the column bars in this and the following figures. ns = not significant for this and the following figures.

### 3.3 Aggressive behavior is reduced in Tg2576 mice treated with MW073 compared to Tg2576 mice treated with vehicle

Next, we determined whether MW073 was capable of reducing the aggressive behavior of Tg2576 male mice. Animals were treated with the 5HT2b receptor antagonist for 3 weeks (daily, i.p.5 mg/kg) prior to performing the aggressivity test. Our data revealed that Tg2576 resident males treated with the serotonin antagonist (n=16) showed less aggressivity than Tg2576 mice treated with vehicle (n=16) during the 10 minutes resident–intruder test sessions (Fig. 3A-D). Specifically, when the intruder was inserted into the cage, resident Tg2576 mice treated with vehicle started the first attack after ∼3 min. By contrast, when the intruder was inserted into the cage, resident Tg2576 mice treated with MW073 started the first attack after ∼4.45 min. Despite the delay in the first attack for Tg mice treated with MW073, the difference was not statistically significant (Fig. 3A). Most importantly, the duration of the first attack in Tg mice treated with MW073 was reduced (Fig. 3B). Moreover, the number of attacks and the time of total attacks decreased in Tg2576 treated with MW073 compared to Tg mice treated with vehicle (Fig. 3C-D).

**Fig. 3.**
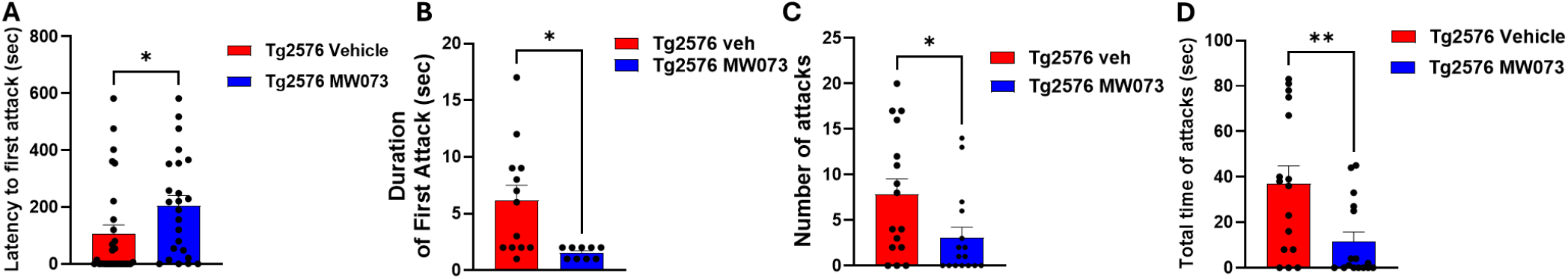
Tg2576 mice treated with MW073 showed amelioration of aggressive behavior. Bar graphs representing rate scores related to aggressive behavior in Tg2576 mice treated with vehicle and Tg2576 treated with MW073. (**A**) Tg2576 resident males treated with MW073 displayed a statistically different latency to the first attack compared to Tg2576 treated with vehicle. (vehicle =198.2 ± 53.44 sec; MW073 = 266.8 ± 45.75 sec, p=0.454) (**B**) Tg2576 resident males treated with MW073 showed a decrease in the duration of the first attack (vehicle = 6.154 ± 1.33 sec; MW073 = 1.556 ± 0.17, p = 0.01). (**C-D**) The number of attacks, as well as the total attack time, of Tg2576 resident males treated with MW073 was reduced compared to Tg2576 treated with vehicle (number of attacks: vehicle = 7.813 ± 1.72, MW073 = 3.063 ± 1.15, p =0.029; total attack time: vehicle = 37.00 ± 7.77, MW073 = 11.56 ± 4.23, p = 0.0074). These experiments were performed on 16 Tg2576 mice either treated with vehicle or MW073. The average age of Tg2576 mice treated with vehicles was 294.56 ± 34.550, whereas the average of mice treated with MW073 was 280.81 ± 32.954 (p = 0.7753).

### 3.4 No differences are evident in control Open Field test in nTg mice, nTg mice treated with MW073, Tg mice treated with vehicle and Tg mice treated with MW073 as well

To exclude the possibility that the beneficial effects of MW073 in Tg2576 mice were secondary to changes in locomotor activity, anxiety-like behavior, or exploratory drive, we assessed animals in the open field test after 3 weeks of daily treatment with the 5-HT2B receptor antagonist (5 mg/kg, i.p.). Analysis of time spent in the center versus the periphery of the arena revealed no significant differences between MW073- and vehicle-treated groups, either in nTg males (n = 14/group) or Tg2576 males (MW073, n = 15; vehicle, n = 16) during the 10-minute sessions on days 1 and 2 (Fig. 4A). As expected, all groups spent more time in the center on day 2 relative to day 1, reflecting reduced anxiety and successful memory-based habituation.

**Fig. 4.**
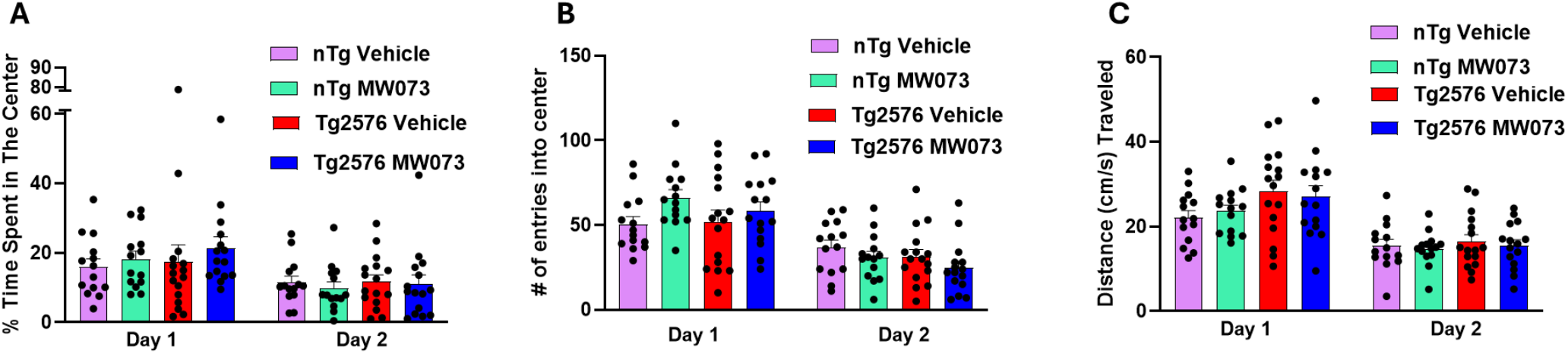
nTg and Tg2576 mice treated with MW073 or vehicle do not show difference in open field control test. Bar graphs representing rate scores related to time spent in the center, number of entries into center and the distance travelled in nTg and Tg2576 mice treated with vehicle and Tg2576 treated with MW073. (**A**) nTg and Tg2576 males treated with MW073 showed no differences compared to nTg and Tg2576 treated with vehicle during the test (day 1: nTg vehicle = 16.34 ± 2.37 sec; nTg MW073 = 18.2 ± 2.27 sec, Tg2576 vehicle = 17.48 ± 4.75 sec; Tg2576 MW073 = 21.41 ± 3.15 sec p=0.7149 ns. Day 2: nTg vehicle = 11.52 ± 1.74 sec; nTg MW073 = 9.83 ± 1.80 sec, Tg2576 vehicle = 11.71 ± 1.92 sec; Tg2576 MW073 = 10.96 ± 2.69 sec p=0.7149 ns.) (**B**) nTg and Tg2576 males treated with MW073 showed no differences compared to nTg and Tg2576 treated with vehicle during the test (day 1: nTg vehicle = 51.62 ± 4.52; nTg MW073 = 66.07 ± 4.84, Tg2576 vehicle = 52.12 ± 6.80; Tg2576 MW073 = 58.34 ± 5.35 p=0.2093 ns. Day 2: nTg vehicle = 36.92 ± 4.21; nTg MW073 = 30.78 ± 3.72, Tg2576 vehicle = 31.37 ± 4.15; Tg2576 MW073 = 24.67 ± 4.02 p=0.2217 ns.) (**C**) nTg and Tg2576 males treated with MW073 showed no differences compared to nTg and Tg2576 treated with vehicle during the test (day 1: nTg vehicle = 22.07 ± 1.66 cm/s; nTg MW073 = 23.63 ± 1.45 cm/s, Tg2576 vehicle = 28.97 ± 2.92 cm/s; Tg2576 MW073 = 27.72 ± 2.63 cm/s p=0.1970 ns. Day 2: nTg vehicle = 15.48 ± 1.51 cm/s; nTg MW073 = 14.78 ± 1.07 cm/s, Tg2576 vehicle = 17.41 ± 1.68 cm/s; Tg2576 MW073 = 15.38 ± 1.51 cm/s p=0.8698 ns.) These experiments were performed on 14 nTg mice either treated with vehicle or MW073, 15 Tg2576 treated with vehicle and 16 Tg2576.

**Fig. 5.**
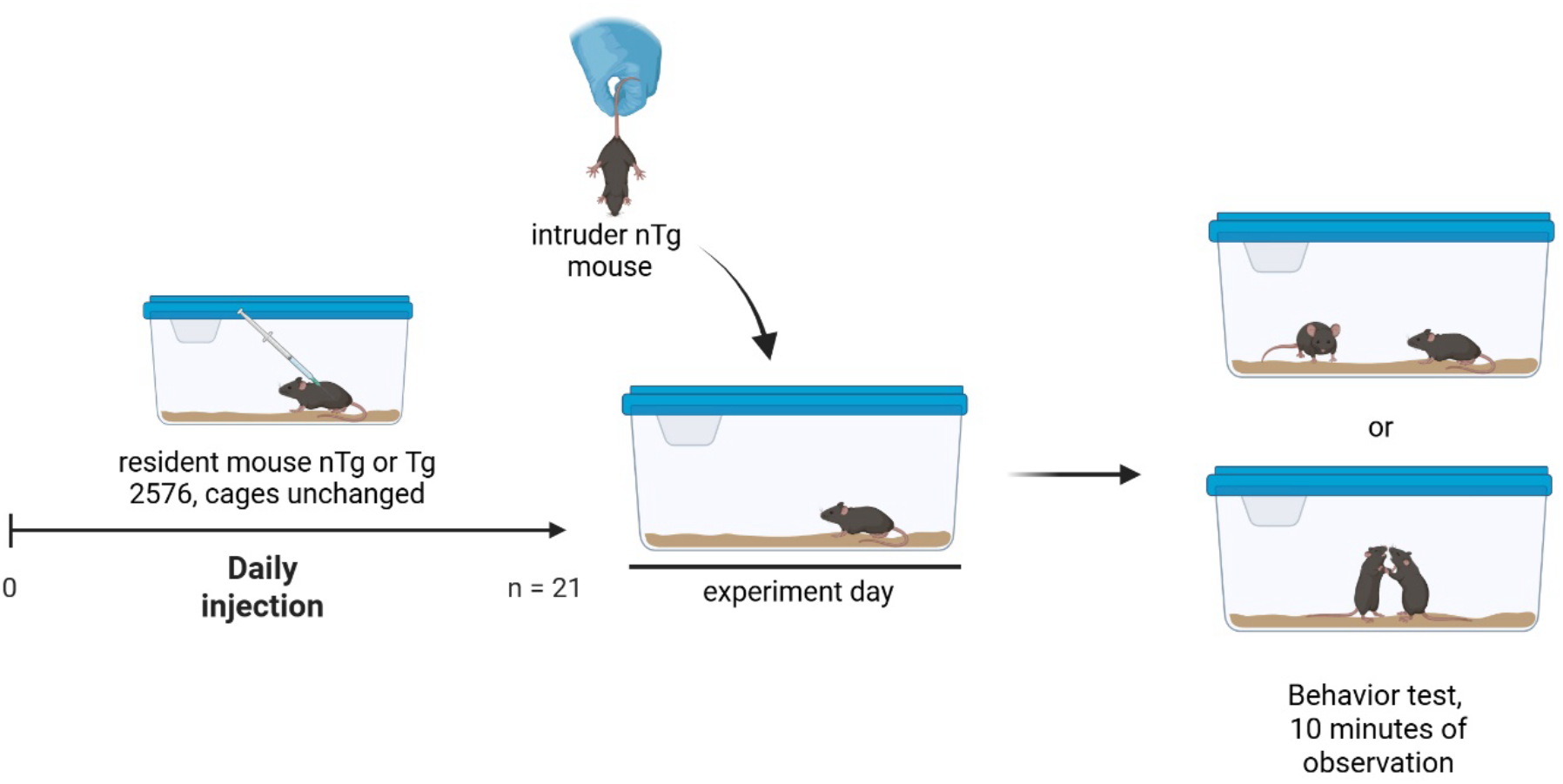
Experimental design. Resident mice (nTg or Tg) are in isolated cage during 21 days of injections. On the test day, an intruder nTg mouse is added in the resident cage and the social behavior animals are observed and recorded for 10 minutes directly after introduction of intruder mice. (Figure created using BioRender (BioRender.com))

Similarly, the number of center entries, another measure of anxiety-like behavior, did not differ between MW073- and vehicle-treated animals in either genotype (Fig. 4B). Again, all groups exhibited increased center entries on day 2, consistent with reduced anxiety and enhanced exploratory drive.

Finally, total distance traveled, a measure of general locomotor activity, was comparable across treatment groups in both nTg and Tg2576 males (Fig. 4C). All groups showed decreased activity on day 2 relative to day 1, consistent with habituation to the test environment. Taken together, these data indicate that MW073 does not alter locomotor function, anxiety-like behavior, or exploratory activity, supporting the conclusion that its effects on aggression in Tg2576 mice are not confounded by changes in these parameters.

## 4. METHODS

### 4.1 Animals and housing conditions

The following groups of mice were used: a) C57BL/6 mice used like control in each experiment, b) Tg2576 mice that overexpress human mutant APP (isoform 695) containing the double mutation K670N, M671L (Swedish mutation) under the control of the hamster prion protein promoter. They are characterized by elevated levels of Aβ and ultimately amyloid plaques. They were obtained from a mouse colony that was established thanks to generous gift of Karen Hsiao-Ashe. Animals were maintained on a 12-hour light/12-hour dark cycle, in a temperature- and humidity-controlled room. Food and water were available ad libitum. Mice were allocated to a specific treatment and paradigm by a randomization procedure. Investigators who performed the experiments were blind in respect to genotype and treatment.

### 4.2 Treatment solution

MW073 compound for treatment in behavioral testing was diluted in 10% Propylene Glycol (Sigma-Aldrich P4347) MilliQ quality water and 0.1% formic acid to prepare a stock solution. At the time of the experiment, the compound was diluted in sterile saline and administered by i.p. injection at concentration of 5mg/kg.

### 4.3 Experimental design

This study aimed to elucidate the effect of antagonist serotonin compound on spontaneous social behavior mice. Mice aggressive behavior will be assessed by means of a combined isolation-induced and resident-intruder paradigm. For this purpose, mice were isolated for 3 weeks. The mice were single housed in their standard cages and left undisturbed during the entire isolation period. Meanwhile, no fresh bedding material was provided to ascertain that the area becomes their own territory and to evoke aggressive behavior upon intrusion by another mouse of the same sex. After 3 weeks of isolation, mice were allowed adaptation to the observation room in their home cage for at least 1 hr prior to testing. A group-housed male C57Bl6 mouse was introduced into the resident’s home cage. Only the behavior of resident mice was analyzed. The second mouse was classified as an intruder. To distinguish the intruder from residence mouse, the intruder was marked with a black sign on the tail. The behavior was recorded for 10 minutes.

### 4.4 Observation of social behavior

During a 10-min observation period, the observer, who was blind to the mouse’s genotype, evaluated animal aggressivity. As an index of social behavior and aggressivity we used mainly pouncing/chasing behaviors (chasing, attacking and escalated fighting) of the mouse discriminating against the approaching, facial/body sniffing, ano-genital sniffing from resident mice. The observer recognized the defensive behavior such as avoiding, fleeing and defensive upright posture.(Kastner et al 2019) The number/severity of physical encounters was closely monitored, and the mice were separated if any encounter was severe enough to potentially cause injury. The number of encounters and latency to encounter were scored using a stopwatch and every test was recorded. (Noldus Information Technology, Wageningen, The Nederland). Also during analysis, the observer was blind to the mice’s genotype treatment.

### 4.5 Open Field control experiment

Spontaneous locomotor activity and anxiety-like behavior were assessed using the Open Field test. Mice were individually placed in the center of a square arena (40 cm × 40 cm, walls 40 cm high) made of white Plexiglas. The test was conducted in a quiet room to minimize external stressors. The animals were allowed to freely explore the arena for 10 minutes. Behavioral activity was recorded using a top-mounted video camera and analyzed with Activity Motor by MedAssociates Inc. The arena was virtually divided into two zones: a central zone (20 × 20 cm) and a peripheral zone. The parameters analyzed included total distance traveled (cm), mean velocity (cm/s), time spent in the center (s), number of entries into the center, rearing behavior, and immobility time. The arena was thoroughly cleaned with 70% ethanol between sessions to eliminate olfactory cues.

To assess habituation, mice were exposed to the same arena for two consecutive days under identical conditions. Changes in locomotor activity and center-related parameters between Day 1 and Day 2 were used as indicators of habituation and exploratory memory. Animals were acclimated to the testing room for at least 30 minutes prior to the session. Behavioral scoring was conducted blind to treatment conditions.

### 4.5 Statistical analysis

Investigators who performed the experiments were blinded with respect to treatment and genotype. The data were scored as a total duration of each behavior performed during the 10-min observation period. The comparisons between Tg males and females, Tg and nTg mice as well as Tg mice treated with vehicle and Tg treated with MW073 on a single variable were carried out using the Student’s t-test (2-tailed) and one way ANOVA by GraphPad Prism10.

## 5. DISCUSSION

5-HT2B receptor was first recognized 50 years ago in the contraction of the gastric fundus from rat stomach. (Vane 1959) The intracellular mechanisms downstream of the 5-HT_2_B receptor is poorly understand. The 5-HT_2_B receptor, a G protein–coupled receptor (GPCR), is expressed in both neurons and astrocytes, where it contributes to serotonergic modulation of brain function. Emerging evidence indicates that 5-HT_2_B receptors, widely expressed in both neurons and glial cells (particularly astrocytes), trigger complex intracellular signaling cascades involving PLC-β activation, intracellular Ca^2+^release, and transactivation of receptor tyrosine kinases such as EGFR and PDGFR, ultimately leading to stimulation of the ERK1/2 and PI3K/Akt pathways. (Li et al 2008, Peng et al 2018, Wang et al 2021). In cultured astrocytes, 5-HT_2_B receptor activation induces intracellular Ca^2+^transients via phospholipase C-β (PLC-β) and inositol trisphosphate (IP_3_) signaling, mobilizing calcium from intracellular stores. This Ca^2+^rise activates protein kinase C (PKC), matrix metalloproteinases (MMPs), and promotes the release of growth factors such as EGF and PDGF. These in turn transactivate EGFR and PDGFR, leading to downstream activation of the Ras–Raf–MEK–ERK and PI3K/Akt pathways. This cascade regulates genes involved in inflammation, metabolism, and synaptic remodeling.(Sanden et al 2000) Subsequent activation of ERK1/2 and PI3K/Akt regulates gene expression of immediate early genes such as c-Fos, enzymes like cPLA_2_, and several metabolic and neurotrophic molecules involved in astrocytic glutamate uptake and glycolysis (Hertz et al 2015). In neurons, 5-HT_2_B receptor activation has been linked to modulation of neuronal excitability and plasticity, although the pathways are less well characterized than in astrocytes. Some evidence suggests neuronal 5-HT_2_B signaling may also engage ERK and Akt pathways, contributing to activity-dependent gene expression and long-term changes in neuronal responsiveness. (Wang et al 2021) Together, these signaling mechanisms place the 5-HT_2_B receptor at the intersection of neuroinflammation, gliotransmission, and synaptic plasticity, highlighting its relevance in neurodegenerative and neuropsychiatric disorders. It is obvious that a serotonin antagonist, like MW073, could have an effect on these signaling pathways, reducing astrocyte reactivity, neuronal hyperexcitability, and pro-aggressive circuit activation.

Serotonin is a key neuromodulator involved in numerous physiological and pathophysiological processes across human organ systems. In the central nervous system, serotonin regulates a broad array of neuropsychological and behavioral functions, including mood, perception, reward, aggression, appetite, memory, attention, and sexual behavior (Berger et al 2009). Several serotonin receptor subtypes are expressed in distinct brain regions and contribute to the fine-tuning of these complex behaviors. Consistent with our findings, a growing body of evidence indicates that the serotonergic system plays a significant role in the regulation of aggression across various behavioral paradigms. Notably, different serotonin receptor subtypes have been already associated with distinct aspects of aggressive behavior. For instance, the 5-HT_1_A and 5-HT_1B_ receptors located in the prefrontal cortex (PFC) have been shown to modulate reactive aggression in the resident–intruder test (Takahashi et al 2012a). Local microinjection of selective 5-HT_1A_ or 5-HT_1_B antagonists (WAY-100635 and SB-224289, respectively) into the PFC significantly alters the frequency and intensity of aggressive episodes, supporting the existence of receptor- and region-specific serotonergic regulation of impulsive aggression. Although MW073 does not target these subtypes, its selective antagonism of the 5-HT_2_B receptor may engage overlapping or interacting circuits. The 5-HT_2_receptor family (5-HT_2_A and 5-HT_2_C) has also been implicated in aggression regulation, though findings vary depending on context and behavioral model. For example, while 5-HT_2_C receptor activation tends to suppress aggression, 5-HT_2_A receptor activation may increase it (Olivier 2004). However, the specific contribution of 5-HT_2_B receptors to aggressive behavior have remained poorly characterized.

Aggression in rodent models is a multifaceted behavioral domain regulated by a complex interplay of neurobiological, genetic, and environmental factors. Accurate assessment of aggressive behaviors is essential for understanding pathophysiological mechanisms and for preclinical evaluation of pharmacological interventions targeting neuropsychiatric or neurodegenerative disorders. Over the years, several behavioral assays have been developed to measure various dimensions of aggression in mice, each with specific strengths and limitations. Among the most employed tests are the Resident-Intruder (RI) test, the Tube Dominance test, and the Neutral Arena test. The RI test is the gold standard for quantifying territorial aggression, particularly in male rodents, due to its ecological validity and sensitivity to pharmacological modulation (Koolhaas et al 2013) In the RI paradigm, a “resident” male mouse housed individually for at least one week is confronted with an unfamiliar “intruder” mouse placed in its home cage. Aggressive behaviors, including latency to first attack, number of attacks, duration of aggressive episodes, and threat postures are recorded and scored using well-validated ethological criteria (Nelson & Trainor 2007, Takahashi & Miczek 2014). This setup mimics naturalistic territorial challenges and reliably elicits robust aggressive responses, especially in strains or models prone to hyperaggression. The Tube Test is a non-violent social dominance assay where two mice enter a narrow tube from opposite ends, and the “winner” is the one who forces the other to retreat. While this method provides information on hierarchical positioning, it is limited by the absence of overt aggression and is less sensitive to anxiolytic or anti-aggressive pharmacological manipulations (Wang et al 2014). The Neutral Arena test introduces two unfamiliar mice in a novel environment. Although this avoids the territorial confound, it results in highly variable aggression responses and often fails to elicit consistent aggressive behavior, making it less suitable for pharmacological studies (Takahashi et al 2012b).

Our choice to utilize the Resident-Intruder test in this study is motivated by its superior sensitivity, ethological relevance, and widespread use in aggression research across models of Alzheimer’s disease and serotonergic dysfunction. The RI test is particularly effective in detecting changes in aggression following pharmacological treatment, as it directly engages neural circuits involved in threat appraisal and territoriality, including the hypothalamus and prefrontal cortex (Ferris et al 1997) (Takahashi et al 2012a). Furthermore, the RI test has been extensively used in mouse models of AD, including the Tg2576 mouse, where it has revealed aberrant aggression associated with amyloid pathology and serotonergic dysregulation (Alexander et al 2011). Since our drug candidate, MW073, is a selective 5-HT2B receptor antagonist, it was critical to use a behavioral assay that reflects the serotonergic modulation of aggression. Previous studies have demonstrated that serotonergic manipulation through 5-HT1A, 5-HT1B, 5-HT2A, and 5-HT2B/C receptors can strongly influence aggression in the RI test (de Boer & Koolhaas 2005). Importantly, the RI test is amenable to repeated testing, enabling within-subject designs and longitudinal assessment of treatment effects. However, it also presents certain limitations. Aggressive behavior can be influenced by prior experience, circadian rhythm, strain differences, and housing conditions. To mitigate these factors, we standardized the housing period (21 days of isolation), used age- and weight-matched intruders, and randomized testing times. Despite its limitations, the resident-intruder paradigm remains the most sensitive and reliable test for detecting pharmacologically-induced changes in aggression, particularly in male mice with altered serotonergic level or neurodegenerative pathology. Our findings using the RI test support the role of 5-HT2B receptor antagonism in reducing aggressive behaviors in Tg2576 mice and provide a robust behavioral endpoint for future translational studies.

In mouse models of AD, transgenic manipulations typically reproduce key pathological hallmarks such as memory loss, impaired synaptic plasticity, and Aβ plaque accumulation. However, replicating the neuropsychiatric and behavioral symptoms of dementia such as mood disturbances and psychosis remains challenging. Interestingly, male Tg2576 mice exhibit markedly elevated levels of aggression compared to non-transgenic controls (Alexander et al 2011), making them a useful model for investigating AD-associated neuropsychiatric symptoms. Tg2576 mice overexpress a human mutant form of amyloid precursor protein (APP) associated with early-onset familial AD. These animals show early cognitive impairments around 5 months of age, including deficits in fear conditioning, and develop significant Aβ plaque deposition by approximately 13 months of age. In our preliminary experiments, we observed that female Tg2576 mice did not display abnormal aggressive behaviors. Therefore, all subsequent behavioral assessments focused exclusively on male mice. Consistent with previous reports (Kastner et al 2019), we found that both male and female mice engaged in social behaviors such as anogenital and nasal sniffing; however, aggressive behaviors were robust and specific to Tg2576 males. In our study, Tg2576 males exhibited significantly greater aggression than nTg controls, characterized not only by a faster initiation of attacks but also by a higher frequency of aggressive episodes. Administration of MW073 led to a reduction in both the number and duration of attacks, suggesting that selective 5-HT_2_B receptor inhibition may be a promising therapeutic strategy to mitigate pathological aggression. Supporting this hypothesis, previous studies have shown that 5-HT_1_A receptor agonists like alnespirone reduce aggression without affecting locomotion or social engagement (de Boer et al 2000). This pharmacological specificity aligns with our observations: MW073 reduced aggressive behavior without inducing sedation or hypoactivity. This distinction is crucial in identifying agents that truly reduce aggression, as opposed to nonspecifically suppressing behavior. These findings point to the therapeutic potential of targeting 5-HT_2_B receptors in managing neuropsychiatric symptoms associated with AD.

In summary, aggression is not simply an outcome of general arousal but is a finely regulated behavioral response governed by distinct serotonergic pathways. Our results position 5-HT_2_B receptor antagonist as a novel and selective mechanism for reducing pathological aggression in AD, warranting further investigation.

## Acknowledge

Cecelia Marie Fatta (for assisting with the behavioral experiments), David Kenneth Stack (for assisting with the behavioral exprriments), Steve Bryan Garcia - Gutierrez (for asssting with the mouse colony maintenence)

